# Integrating large-scale meta-GWAS and PigGTEx resources to decipher the genetic basis of complex traits in pig

**DOI:** 10.1101/2023.10.09.561393

**Authors:** Zhiting Xu, Qing Lin, Xiaodian Cai, Zhanming Zhong, Bingjie Li, Jinyan Teng, Haonan Zeng, Yahui Gao, Zexi Cai, Xiaoqing Wang, Liangyu Shi, Xue Wang, Yi Wang, Zipeng Zhang, Yu Lin, Shuli Liu, Hongwei Yin, Zhonghao Bai, Chen Wei, Jun Zhou, Wenjing Zhang, Xiaoke Zhang, Shaolei Shi, Jun Wu, Shuqi Diao, Yuqiang Liu, Xiangchun Pan, Xueyan Feng, Ruiqi Liu, Zhanqin Su, Chengjie Chang, Qianghui Zhu, Yuwei Wu, The PigGTEx Consortium, Zhongyin Zhou, Lijing Bai, Kui Li, Qishan Wang, Yuchun Pan, Zhong Xu, Xianwen Peng, Shuqi Mei, Delin Mo, Xiaohong Liu, Hao Zhang, Xiaolong Yuan, Yang Liu, George E. Liu, Guosheng Su, Goutam Sahana, Mogens Sandø Lund, Li Ma, Ruidong Xiang, Xia Shen, Pinghua Li, Ruihuang Huang, Maria Ballester, Daniel Crespo-Piazuelo, Marcel Amills, Alex Clop, Peter Karlskov-Mortensen, Merete Fredholm, Guoqing Tang, Mingzhou Li, Xuewei Li, Xiangdong Ding, Jiaqi Li, Yaosheng Chen, Qin Zhang, Yunxiang Zhao, Fuping Zhao, Lingzhao Fang, Zhe Zhang

## Abstract

Understanding the molecular and cellular mechanisms that underlie complex traits in pigs is crucial for enhancing their genetic improvement program and unleashing their substantial potentials in human biomedicine research. Here, we conducted a meta-GWAS analysis for 232 complex traits with 28.3 million imputed whole-genome sequence variants in 70,328 individuals from 14 pig breeds. We identified a total of 6,878 genomic regions associated with 139 complex traits. By integrating with the Pig Genotype-Tissue Expression (PigGTEx) resource, we systemically explored the biological context and regulatory circuits through which these trait-associated variants act and finally prioritized 16,664 variant-gene-tissue-trait circuits. For instance, rs344053754 regulates the expression of *UGT2B31* in the liver by affecting the activity of regulatory elements and ultimately influences litter weight at weaning. Furthermore, we investigated the conservation of genetic and regulatory mechanisms underlying 136 human traits and 232 pig traits. Overall, our multi-breed meta-GWAS in pigs provides invaluable resources and novel insights for understanding the regulatory and evolutionary mechanisms of complex traits in both pigs and humans.

## Introduction

Pigs are globally recognized as one of the most important farm animals, with pork production reaching 106.1 million tons in 2021^1^. Understanding the genetic control of complex phenotypes in pigs help us genetically maximize their production efficiency^2^, and improve their health and welfare^3,4^, while minimizing environmental challenges^5,6^ through advanced precision breeding techniques. For example, genomic selection substantially and durably enhance the efficiency of pig breeding programs in terms of reliability, genetic trends, and inbreeding rates^7^. Genome editing also protect pigs from porcine reproductive and respiratory syndrome virus and reduce economic losses^8^. On top of their great economic importance as a primary source of animal protein for humans^9^, pigs have been widely accepted as a model for studying human biology and diseases, including Alzheimer’s disease^10^, cardiovascular disease^11^, wound healing^12,13^, human reproduction^14^, the human gastrointestinal tract^15^, dry eye^16^, and immunological studies^17–19^. Therefore, investigating the genetic and biological architecture of complex traits in pigs will not only benefit the pig breeding industry but also to human biomedical research.

Performing a genome-wide association study (GWAS) is a commonly used strategy for dissecting complex trait/disease genetics^20–22^. As of June 10, 2023, the Pig quantitative trait loci (QTL) database (Pig QTLdb) has reported 48,844 QTL, representing 673 distinct traits and 279 trait variants^23^. However, causal variants and genes underlying most of these QTL regions remain unknown due to the large amount of linkage disequilibrium (LD) of genetic variants within pig populations/breeds^24^. Cross-ancestry/population meta-GWAS analysis has been proposed as an efficient approach for identifying trait-associated variants shared between populations and accelerate statistical fine-mapping of causal variants and genes through reducing LD^22,25–27^. In addition, the majority of genetic variants identified in GWAS were located in non-coding genomic regions^28^, and were significantly enriched in *cis*-regulatory elements, including promoters and enhancers^29,30^, as well as gene expression QTL (eQTL) in relevant tissues^31^. This suggests that GWAS variants might exert their effect *via* regulating gene expression. Therefore, it is of great interest to prioritize the causal variants, genes, pathways, and tissues of complex traits through systematically integrating functional annotation data such as FAANG^32^ and FarmGTEx resources^33^.

Here, we collected and analyzed phenotypes and genotypes of 70,328 pigs from 59 populations representing 14 pig breeds to identify genetic variants underlying complex traits in pigs. After imputing genotypes to a whole genome sequence level using a multi-breed reference panel^33^, we conducted a comprehensive cross-population/breed meta-GWAS analysis for 232 complex traits. We then integrated multi-tissue regulatory elements from the FAANG project^32^ and multi-tissue molQTL from the PigGTEx project^33^ to systematically resolve the functional molecular basis of complex traits in pigs. To further investigate the potential of pigs as model organisms for human biology and diseases, we compared the genetic regulations of 136 human complex phenotypes and 232 pig complex phenotypes. Finally, we developed an open-access and user-friendly website (http://pigbiobank.ipiginc.com/home) for the research community to query and download the genetic associations of complex traits in pigs (Fig. 1a).

**Figure 1.**
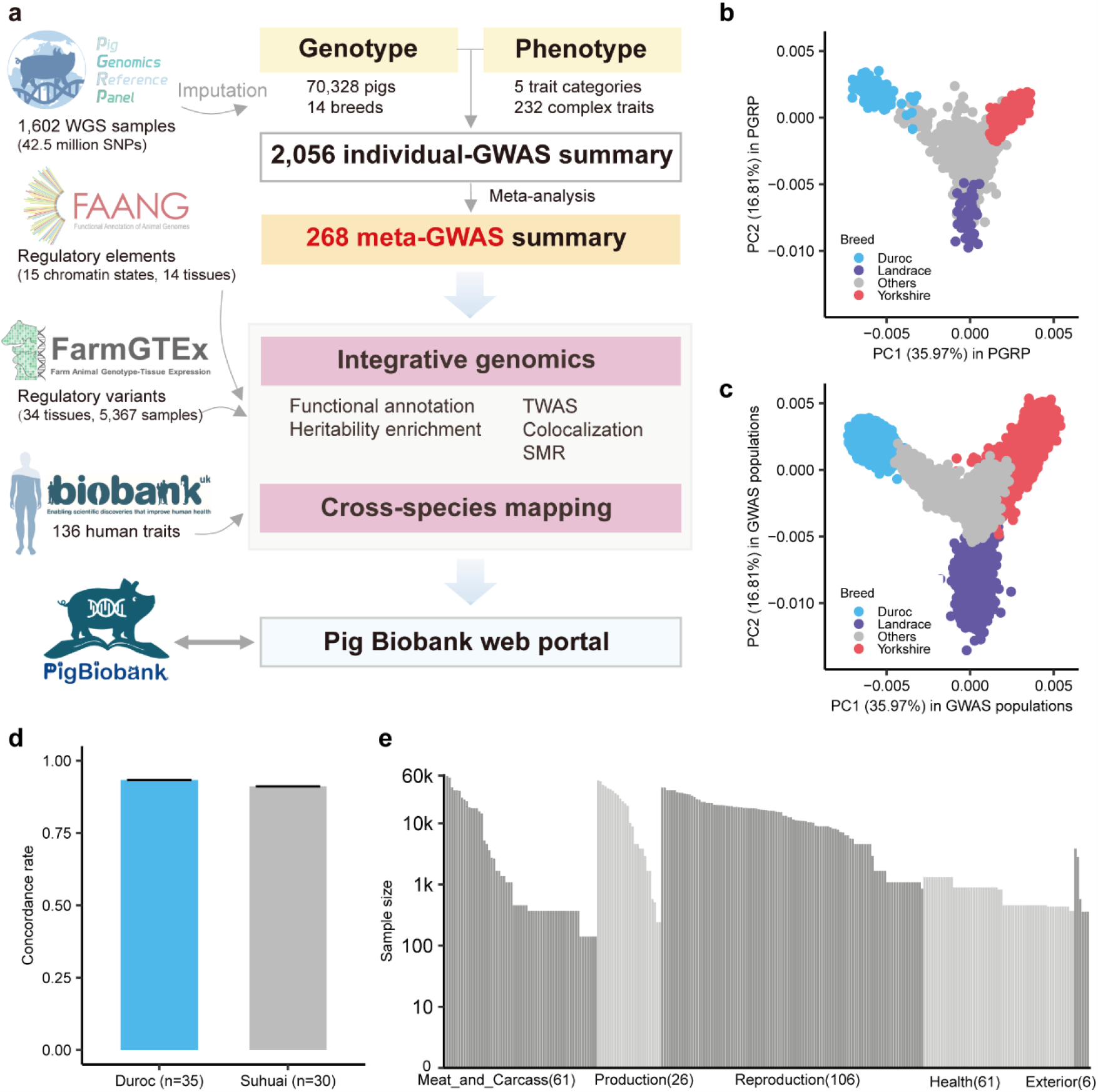
The overall study design and summary of genotypes and phenotypes. **(a)** Overview of study design. Genotyping arrays: Illumina Porcine SNP60K Bead Chip (N = 10,870), the GeneSeek Genomic Profiler (GGP) Porcine SNP80 BeadChip (N = 4,724), the GGP Procine SNP50 BeadChip (N = 29,789), the KPS Porcine Breeding Chip (N = 21,618), the GenoBaits Porcine SNP50K BeadChip (N = 454) and low-coverage sequence (N = 2,873). WGS: Whole genome sequence. GWAS: genome-wide association study. TWAS: transcriptome-wide association study. SMR: summary data-based Mendelian randomization. **(b-c)** Principal component analysis of PGRP **(b)** and GWAS **(c)** populations, which were conducted based on all 57,600 individuals (samples with genotype data) and a total of 1,603 shared array SNPs using PLINK (v1.90)^34^ (parameters: --geno 0.1 --mind 0.1 --indep-pairwise 50 5 0.5 --maf 0.01 and --pca 10). The first two principal components were plotted using the geom_point function from ggplot2 (v3.3.6) in R (v4.1.2). **(d)** The imputation accuracy of PGRP in independent WGS data. This was 93.34% ± 7.64% for Duroc pigs (commercial breed and within PGRP) and 91.13% ± 10.49% for Suhuai pigs (domesticated breed and outside PGRP). The imputation accuracy was calculated as the concordance rate between the imputed and observed genotypes. **(e)** The total sample size for each trait in meta-GWAS analyses. Traits were classified into five main categories.

## Results

### Summary of genotypes and phenotypes

After excluding ancestral outliers within each of the 59 populations based on population structure analysis (details see Methods), we retained 69,242 pigs genotyped by SNP arrays (with an average of 45,418 autosomal SNPs) or low-coverage whole-genome sequencing (WGS) (with 198,178 autosomal SNPs) for subsequent analysis, including 20,706 Duroc, 9,159 Landrace, 34,540 Yorkshire, and 4,837 individuals from 11 other breeds (Table S1). We imputed genotypes of all the 69,242 animals to the sequence level, using the multi-breed **P**ig **G**enomics **R**eference **P**anel (PGRP version 1) from the PigGTEx project^33^ as the reference panel, which comprises 42,523,218 autosomal biallelic SNPs from 1,602 WGS data worldwide (Fig. 1b-c). The average concordance rates and genotype correlations between imputed and observed genotypes were 96.67% and 93.86%, respectively, across breeds (Fig. S1a-c, Table S1). We further assessed the genotype imputation accuracy in 65 WGS samples (35 Duroc and 30 Suhuai) that were independent of PGRP (Table S2). The observed concordance rates between the imputed and WGS-called genotypes were 93.34% and 91.13% (genotype correlations of 90.21% and 87.22%), respectively (Fig. 1d, S1d). The genotype imputation accuracy was influenced by the minor allele frequency (MAF) and dosage R-squared (DR^2^, the estimated squared correlation between the estimated allele dose and the true allele dose) (Fig. S1e-h). We thus considered 28,297,602 SNPs with both DR^2^ > 0.8 and MAF ≥ 0.01 in each population for subsequent analysis. As expected, the population structure of GWAS samples estimated by imputed genotypes was consistent with that estimated from raw genotypes (correlation > 0.99) (Fig. S1i). The imputed SNPs were evenly distributed across diverse genomic features (Fig. S1j-l). Altogether, these results supported the reliability of our imputed genotype data.

In total, we collected 271 continuous traits across 59 pig populations in 14 breeds, with an average sample size of 1,141 for each population and each trait (ranging from 116 in Total number of born to 9,246 in Average daily gain), representing 5 main trait categories and 17 subcategories: **Production** (n = 57,612; Feed intake (n = 240), Growth (n = 57,612), Feed conversion (n = 19,095)), **Meat and Carcass** (n = 65,883; Fatness (n = 60,203), Anatomy (n = 52,470), Chemistry (n = 368), Fatty acid content (n = 368), Texture (n = 140), Meat color (n = 140), pH (n = 140)), **Health** (n = 2,139; Immune capacity (n = 1,317), Blood parameters (n = 2,139)), **Reproduction** (n = 71,637; Reproductive traits (n = 41,569), Litter traits (n = 51,717), Reproductive organs (n = 40,914)), and **Exterior** (n = 6,625; Behavioral (n = 2,797), Conformation (n = 3,828)) (Fig. S2a). In addition, we collected 15 binary traits across 23 populations in 3 breeds with an average sample size of 1,025 (ranging from 160 in Number of mummified pigs of parity 1 to 9,246 in Teat number symmetry), representing Reproduction category (Litter traits (n = 12,655) and Reproductive organs (n = 24,087)) (Fig. S2a). After filtering out samples with low-quality genotypes and phenotypes (Methods), we retained 249 continuous traits (average sample size of 1,136) and 11 binary traits (average sample size of 1,035) for subsequent analysis. The average backfat thickness (M_BFT) had the largest cumulative sample size of 58,725 (Fig. 1e). Across all the traits, we observed an average heritability of 0.27, ranging from 0.02 for Teat number (left) to 0.97 for Lysozyme level (Table S3, Fig. S2b-c).

### Individual GWAS and meta-GWAS analysis

We conducted GWAS for 249 individual continuous traits and 11 binary traits in each population, yielding a total of 2,117 GWAS summary statistics (Table S3). To ensure the quality and reliability of these individual GWAS results for subsequent meta-analysis, we applied stringent quality control using multiple strategies, including SE-N plot, P-Z plot, EAF plot, and λ_GC_ (Methods). This resulted in 2,056 high-quality GWAS summary statistics, representing 221 continuous traits and 11 binary traits (Fig. S2d-j). Of these, 78 traits were not previously reported in Pig QTLdb (release 46)^23^ (https://www.animalgenome.org/cgi-bin/QTLdb/SS/index). In total, we detected 8,098 QTLs (*P* < 5×10^−8^) for 154 traits, representing 7,011 non-overlapping lead SNPs (5,665 SNPs with m-value > 0.9 in at least one study, while the m-value represents the posterior probability of the effect estimated by METASOFT). The correlations of SNP effects were significantly higher for the same traits across different populations/breeds compared to different traits within the same populations/breeds (Fig. S3a-b). Interestingly, among the 5,665 lead SNPs, 69.88% were only detected in one population for a specific trait (Fig. S3c), and their MAFs were higher in the target populations compared to the remaining populations (Fig. S3d). For instance, *rs323720776* was associated with Average daily gain (birth-100kg) only in a Yorkshire pig population, with its MAF in this population (MAF = 0.46) being higher than in others (average MAF = 0.27) (Fig. S3e). These findings suggest that population-specific associations may arise from differences in variant segregation between populations.

To detect population-shared associations with small effect sizes that could not be detected by individual GWAS due to limited sample size^35^, we conducted meta-GWAS analyses for each of the 232 complex traits across populations/breeds using individual GWAS summary statistics. Out of these traits, 25 common traits had larger sample sizes and were classified as main traits (M_traits), covering the categories of Growth, Fatness, Reproductive, Anatomy, Reproductive organs and Litter trait (Table S3). Furthermore, we conducted 36 within-breed meta-GWAS analyses for the Duroc, Landrace and Yorkshire breeds, focusing on 12 M_traits with large sample sizes in all of the three breeds (prefixed with ‘D_’, ‘L_’ and ‘Y_’, respectively), to explore potential breed-specific genetic regulation mechanisms for complex traits. The average sample size of these 268 meta-GWAS analyses was 6,409, ranging from 137 for dressing percentage to 56,165 for M_BFT (Table S4). Overall, we detected 6,878 QTLs for 139 traits in 169 meta-GWAS analyses (*P* < 5×10^−8^), representing 6,233 non-overlapping lead SNPs (Table S5, Fig. 2a). These lead SNPs were distributed across all the 18 autosomes (Fig. 2b) and had smaller MAFs than random SNPs (Fig. S4a). Furthermore, the number of significant QTLs detected in meta-GWAS showed positive correlations with both sample size (Pearson’s *r* = 0.69, *P* = 1.36×10^−25^) and trait heritability (Pearson’s *r* = 0.49, *P* = 9.59×10^−3^) (Fig. 2c-d), consistent with findings in humans^36,37^.

**Figure 2.**
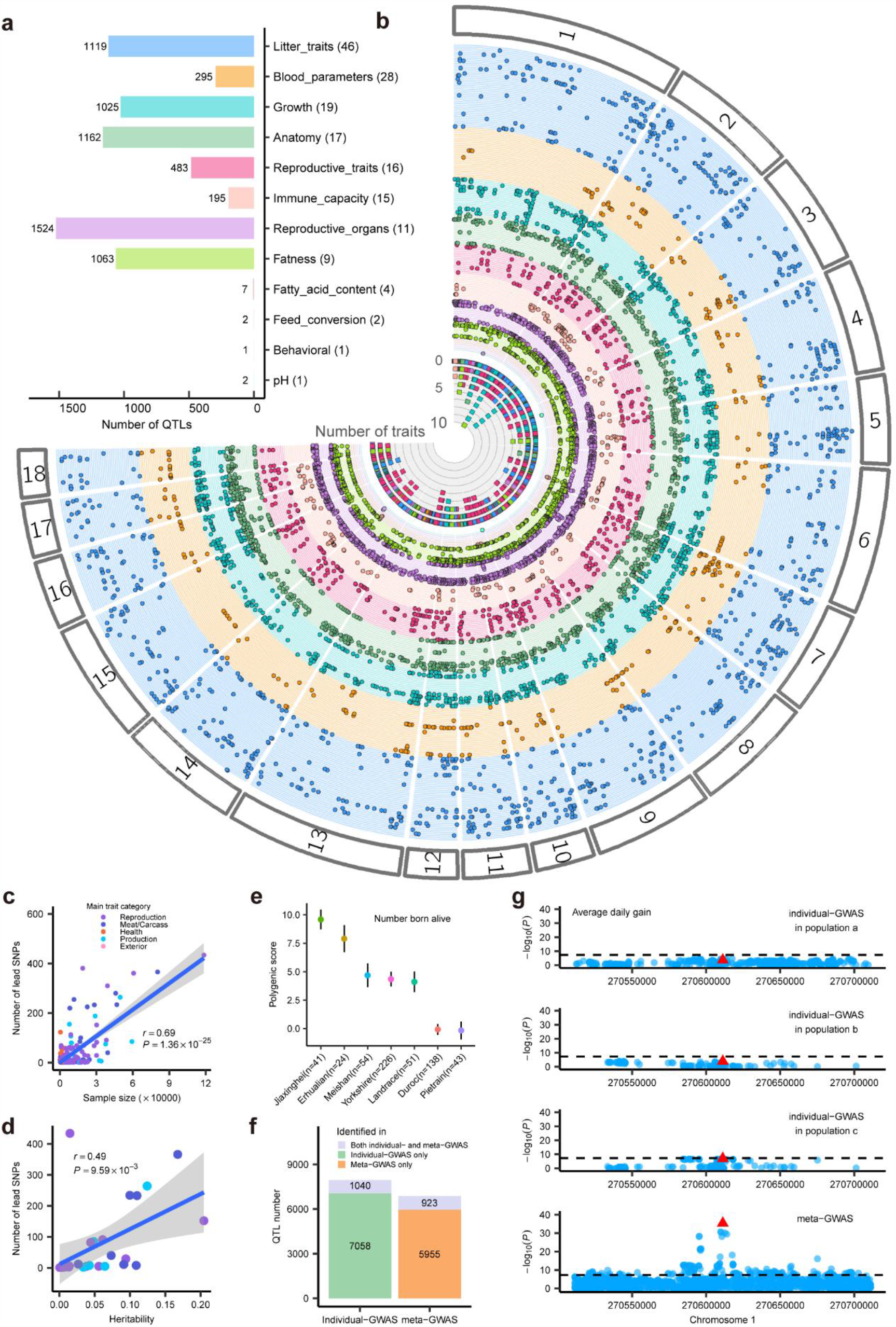
Summary and validation of quantitative trait loci (QTL) for pig complex traits. **(a)** The number of QTL for 12 sub trait-categories. **(b)** Fuji-plot summarizes the 6,878 lead SNPs (*P* < 5×10^−8^) identified in 169 meta-GWAS analyses. It was completed using the Fuji-plot script developed by Kanai et al.^42^ The inner-most ring (ring 1) indicates the number of traits associated with each SNP. Rings 2-170 indicate the 169 traits. The order of traits is shown in Table S3 (starting with the inner-most ring). The points indicate the genomic position of the 6,878 SNPs associated with the traits. **(c)** Pearson correlation between sample size and the number of lead SNPs (*P* < 5× 10^−8^) in 169 meta-GWASs with lead SNPs detected. **(d)** Pearson correlation between heritability and the number of lead SNPs (*P* < 5×10^−8^) in 27 meta-GWASs (sample size > 15,000). Heritability was estimated using LD score regression (LDSC)^43^. The Pearson correlation coefficient in **(c-d)** was calculated by the *cor*.*test* function in R. **(e)** Results of genomic predictions for individuals from several pig breeds in the PGRP with large phenotype differences, based on a linear mixed model and genomic information from suggestive lead variants (*P* < 1×10^−5^) in the total number born alive (M_NBA). The x-axis labels indicate the different pig breeds. The y-axis labels indicate the genomic estimated breeding values (GEBVs). The black error bars are the standard errors of GEBVs. **(f)** The number of different categories of QTLs detected in individual GWAS and meta-GWAS. **(g)** rs320375241 associated with Average daily gain (M_ADG) in individual GWASs of population a, b, and c and meta GWAS, respectively. The a, b, and c were three random populations for M_ADG.

In comparison to Pig QTLdb (release 46)^23^, we identified 14,704 novel QTLs for 209 traits in both individual GWAS and meta-GWAS (Fig. S4b-c). Furthermore, we employed two strategies to validate the detected lead SNPs. Firstly, we conducted meta-GWAS for average daily gain (ADG) in independent populations, including 42,790 Duroc pigs, 88,984 Landrace pigs, and 69,606 Yorkshire pigs. The association signals detected in these different independent populations were significantly enriched in the QTL regions of ADG detected in this study (Fig. S4d). Secondly, we used suggestively significant lead SNPs (*P* < 1×10^−5^) to predict genetic values of litter size/teat number across seven pig breeds. We observed that Jiaxinghei, Erhualian, and Meishan pigs exhibited higher predicted values than Landrace and Duroc (Fig. 2e, Fig. S4e). This aligns with previous findings that Meishan pigs maintained a higher number of follicles during the follicular phase than Landrace hybrid pigs^38^. In summary, these results illustrate that lead SNPs detected here are reliable and shared among populations/breeds.

In comparison with individual GWASs, we identified 5,955 novel QTLs in meta-analyses for 147 traits (Fig. 2f). For instance, *rs320375241* was non-significant for ADG in any individual GWASs, but was identified as a significant lead SNP of ADG in the meta-analysis (Fig. 2g). Furthermore, we found 7,058 QTLs in individual GWASs that were not detected in the meta-GWASs (referred to as class A QTLs) (Fig. 2f). When compared to the QTLs detected in both individual GWASs and meta-GWASs (referred to as class B QTLs), the lead SNPs of class A QTLs tended to have different directions of effects on the trait across study populations (Fig. S4f). Additionally, the populations in which class A QTLs were detected had a smaller proportion of the total sample size in the meta-analysis of the trait (Fig. S4g). These findings suggest that GWAS variants with opposite directions of effect among populations, or GWAS variants detected in populations with small sample sizes, may result in undetectable QTLs in meta-analyses.

To characterize the genetic regulation of complex traits (Fig. S3c), we here only considered QTLs/lead SNPs identified in the meta-GWAS analysis. Across all the 64 meta-analyses with a number of lead SNPs > 10, we observed a negative correlation between MAF and effect size of lead SNPs, with a median Pearson correlation of -0.65, ranging from -0.88 in the GGT trait to -0.27 in the number of mummified pigs (Fig. S5a). This suggests that variants significantly associated with complex traits might be under negative selection, similar to previous findings in humans^39–41^. Among these correlations, we observed differences among Duroc, Landrace and Yorkshire in the correlations between MAF and effect size of lead SNPs for ADG and teat number (TNUM). Specifically, the negative correlation for ADG was weaker in Duroc compared to Landrace and Yorkshire (Fig. S5b), while the ADG phenotype value in Duroc was higher than in Landrace and Yorkshire (Fig. S5c). Of note, we found the opposite result for TNUM (Fig. S5d-e). This finding suggests that the three breeds have undergone different levels of artificial selection for different complex traits of economic importance. In the all-breed meta-analyses of 12 M_traits, we identified 1,460 novel QTLs compared to within-breed meta-analyses in Duroc, Landrace, and Yorkshire (Fig. S5f). Most of the breed-specific QTLs (median 77%) detected in the within-breed meta-analysis exhibited different directions of effect between breeds (Fig. S5g). For instance, the effect size of the lead SNP 6_163238739_A_G for ADG in Duroc was 7.53 (*P* = 3.22×10^−8^), while it was -4.40 (*P* = 0.18) in Landrace and -3.99 (*P* = 0.05) in Yorkshire, while this SNP was not associated with ADG in the all-breed meta-analysis (effect size = 2.58, *P* = 0.01) (Fig. S5h). This suggests that associations with different directions of effect between breeds will be offset in the all-breed meta-GWAS analysis, leading to reduced statistical power.

### Pleiotropy of genetic variants in complex traits

To explore breed-specific and shared trait-associations among Duroc, Landrace and Yorkshire breeds, we estimated the posterior probability (m-value) of lead SNPs for each trait using METASOFT^44^. In 12 M_traits, we identified 6,624 SNPs exclusively in one breed and 2,378 SNPs in at least two breeds (m-value > 0.9). For example, in BFT, 1,840 SNPs were exclusively detected in one breed, while 667 SNPs were detected in at least two breeds (Fig. 3a). Breed-specific trait-associated variants had higher MAFs in the breed where they were detected compared to the other breeds (Fig. 3b). In addition, SNPs exclusively detected in one breed had a significantly greater effect on traits compared to those detected in multiple breeds (Fig. 3c). To gain further insights into the regulatory mechanisms of these breed-specific and shared SNPs, we conducted functional annotation and enrichment analysis. Our results revealed that SNPs detected in all three breeds were significantly enriched in tissue-specific gene regions (less than five tissues) (Fig. 3d). SNPs detected in at least two breeds showed a significantly higher enrichment in active promoters and enhancers compared to breed-specific SNPs (Fig. 3e). Additionally, by examining Z-scores of lead SNPs detected exclusively in one breed, we were able to cluster the 36 meta-GWASs of the 12 M_traits based on breed, whereas by examining Z-scores of lead SNPs detected in all three breeds, we were able to cluster the 36 meta-GWASs of the 12 M_traits based on trait (Fig. S6).

**Figure 3.**
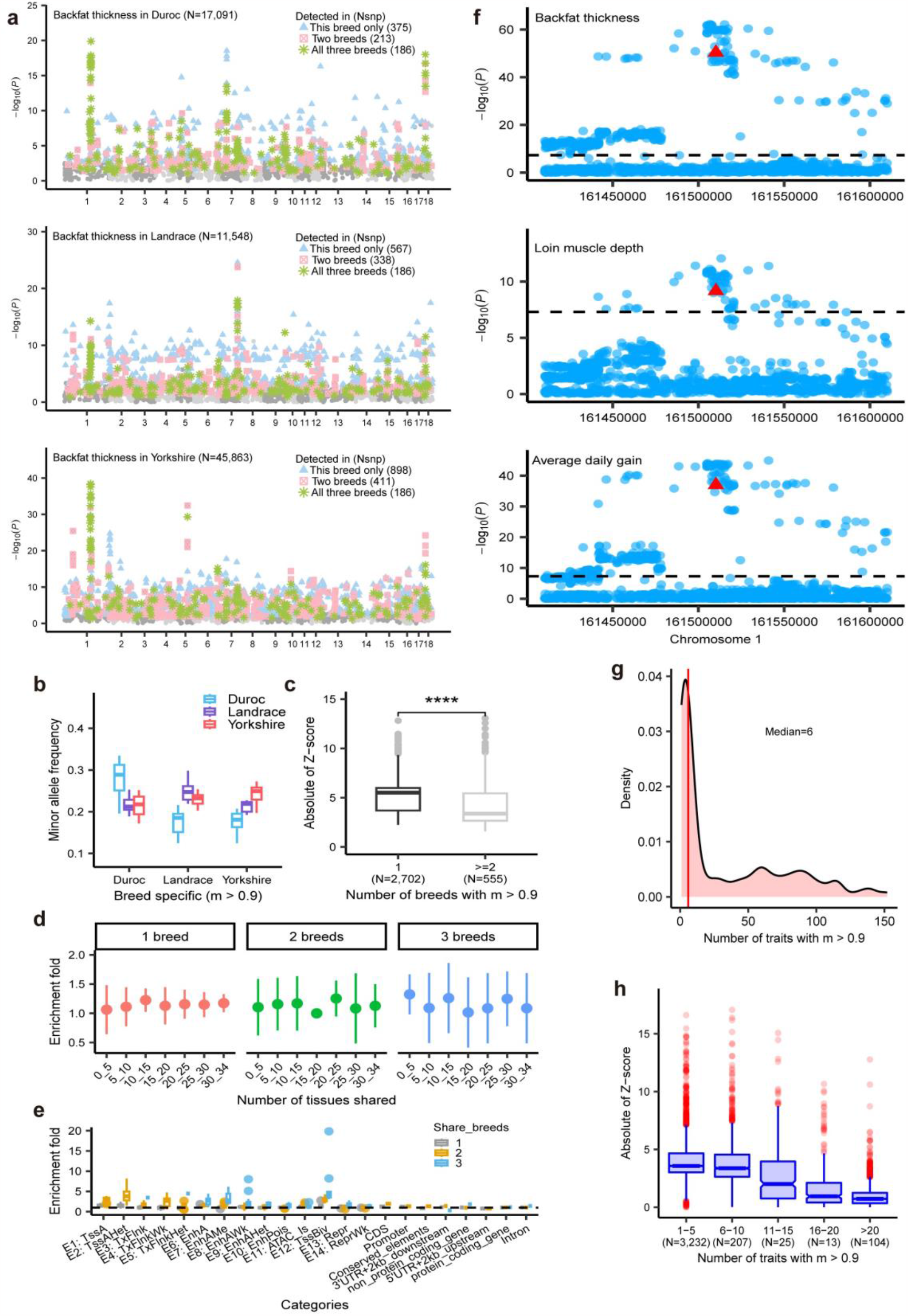
Distribution and functional annotation of QTLs. **(a)**Manhattan plots of backfat thickness (BFT) meta-analysis in Duroc (top), Landrace (middle) and Yorkshire (bottom). Colors and shapes indicate the breed specificity of SNPs with m-values > 0.9. **(b)** Distribution of MAFs in three breeds of traits-associated SNPs detected only in the current breed. **(c)** Distribution of the absolute values of the z-score of SNPs associated with traits detected in different numbers of breeds on traits. The significance of differences between groups was calculated by the *t*.*test* function in R. **(d)** Enrichment of trait-associated SNPs detected in different numbers of breeds in gene regions with different degrees of tissue sharing. **(e)** Enrichment of trait-associated SNPs detected in different numbers of breeds in different categories of genomic regions, conserved elements and regulatory elements. **(f)** Local Manhattan of meta-analysis of M_BFT (top), Loin muscle depth (M_LMDEP) (middle), and M_ADG (bottom) on chromosome 1. **(g)** Density plot of the number of traits for which SNPs associated (m-value > 0.9). **(h)** Distribution of the absolute values of the z-score of SNPs associated with different numbers of traits (m-value > 0.9).

Among 3,581 lead SNPs of 232 traits, 2,100 were associated with at least two traits, with one SNP associated with up to 152 traits (15_21820815_A_G, m-value > 0.9, phastCons = 0.554) (Fig. 3g). For instance, we identified *rs320916522* near *MC4R* on chromosome 1 as being associated with M_ADG (*P* = 1.09×10^−37^, M = 1), M_BFT (*P* = 4.56×10^−51^, M = 1)), and M_LMDEP (*P* = 6.90×10^−10^, M = 1)) (Fig. 3f). *MC4R* has been extensively demonstrated to be linked to muscle and fat deposition in pigs^45–47^. Notably, similar traits shared a greater number of lead SNPs, such as TNUM-related traits (Fig. S7). Moreover, we observed a trend where lead SNPs with higher pleiotropy exhibited smaller effects on traits (Fig. 3h). This result is consistent with the ‘network pleiotropy’ hypothesis proposed by Boyle, Li, and Pritchard, which suggests that small perturbations in a densely connected functional network have at least a small effect on all phenotypes affected by the network^48^.

### Regulatory architecture underlying complex traits

To explore the biological context and regulatory circuits by which the detected trait-associated variants act, we examined multi-layered biological data, including genomic variants, mammalian conserved elements and regulatory elements, to annotate lead SNPs and genome-wide significant SNPs of all the 169 meta-GWASs. Among 6,233 lead SNPs, 0.95% were located in coding regions, while 99.05% were in noncoding regions. Specifically, 43.26% were located in introns, 39.42% in intergenic regions, 9.29% in promoters, and 52.99% in enhancers (Fig. 4a and Fig. S8a). We obtained similar results when analyzing human GWAS variants (Fig. S8b). Lead SNPs were observed to be more concentrated around the transcription start sites (TSS) of protein-coding genes compared to non-lead SNPs (Fig. S8c). Furthermore, lead SNPs exhibited significant enrichment in protein-coding regions (CDS) (18.10-fold, *P* < 1×10^−30^), conserved elements (6.52-fold, *P* = 5.65×10^−8^), and regulatory elements, particularly in active regulatory elements such as active promoters (TssA) and enhancers (EnhA) (Fig. 4a and Fig. S8d-e). Additionally, lead SNPs had lower PhastCons scores (indicating weaker evolutionary constraints) with a median of 0.023, compared to non-lead SNPs (median of 0.062) with matching MAF and linkage disequilibrium (LD) of significant SNPs (Fig. 4b).

**Figure 4.**
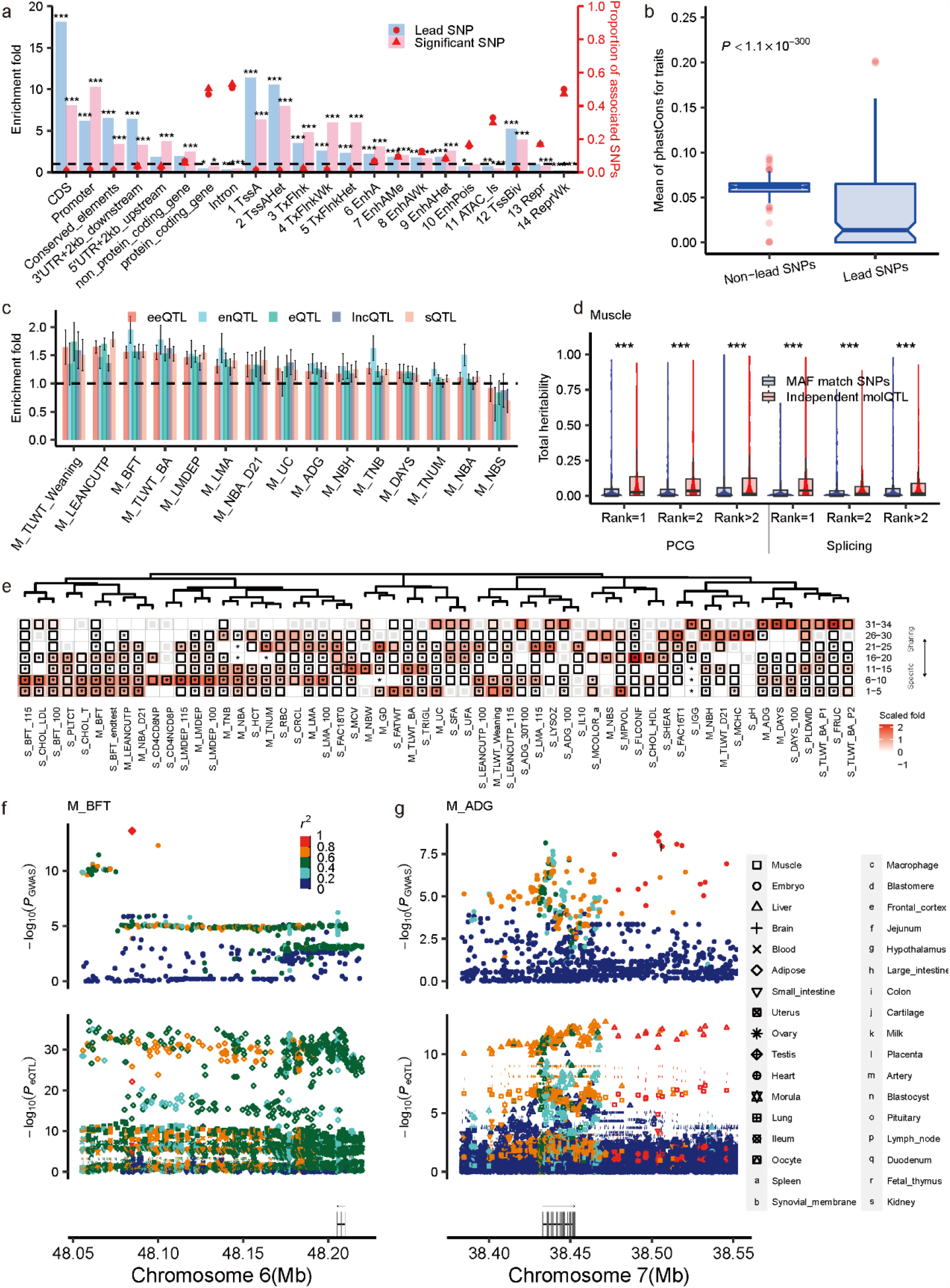
Exploiting the PigGTEx resource to decipher regulatory mechanisms of GWAS loci. **(a)** Results of annotation and enrichment of lead SNP and genome-wide level significant SNPs in different categories of genomic regions, conserved elements and regulatory elements. The red dots indicate the proportion of associated SNPs located in category *C*. The bars indicate the enrichment for category *C*. Significance was indicated by *, ** and *** for *P* < 0.05, 0.01 and 0.001, respectively. **(b)** The mean DNA sequence constraints (PhastCons scores of 100 vertebrate genomes) for lead SNPs in each trait and non-lead SNPs matched with lead SNPs for linkage disequilibrium (within 0.1) and minor allele frequency (MAF) (within 0.02). The *ks*.*test* function of R (v4.1.2) is used to test the difference between groups. **(c)** The heritability enrichment for five types of molecular *cis*-QTLs in 18 main traits. The dashed line represented the enrichment fold = 1. The error bar represented the standard error of the enrichment fold. *cis*-eQTL: gene expression QTL, *cis*-sQTL: splicing QTL, *cis*-eeQTL: exon expression, *cis*-lncQTL: lncRNA expression QTL, *cis*-enQTL: enhancer expression QTL. The details of trait names are described in Table S4. **(d)** The estimated total SNPs heritability contributed by different ranks of independent molecular QTL (molQTL) in Muscle for 268 complex traits. Rank=1, 2 and >2 represented the first, secondary and more than secondary independent molQTLs, respectively. Significance was indicated by *** for *P* < 0.001, which was obtained by the Wilcox test. **(e)** The heritability enrichment for the genes of seven tissue-sharing gradients in 59 complex traits. The red color represented the scaled heritability enrichment fold. Black borders indicated heritability enrichment fold greater than 1. The “*” indicated heritability significant enrichment (Normal test, *P* < 0.05). Column clusters were produced by the *dist* function with the “euclidean” method and the *hclust* function with the “complete” method in R. The heatmap was plotted by ggplot2 package (v3.3.2) in R (v4.2.1). **(f)** The lead SNP rs1108824455 in backfat thickness (M_BFT) was also eQTL of *LGALS13* (tissue-specific magnitude = 10) in five tissues. The top local Manhattan plot was the GWAS of M_BFT for the lead variant (rs1108824455). The middle local Manhattan plot was the eQTL mapping of *LGALS13* for all of the tissues. **(g)** The lead SNP rs324200444 in average daily gain (M_ADG) was also eQTL of *ABCC10* (tissue-sharing magnitude = 33) in four tissues. The top local Manhattan plot was the GWAS of M_ADG for the lead variant (rs324200444). The middle local Manhattan plot was the eQTL mapping of *ABCC10* for all of the tissues. Shapes in **(f-g)** indicate different tissues. The filled colors in **(f-g)** represent linkage disequilibrium. The diagram below in **(f-g)** indicates the positions and strand direction of genes in the locus.

To further investigate the regulatory role of genetic variants on complex traits, we integrated five types of molecular QTLs (molQTLs, including cis-eQTLs for PCG expression, cis-eeQTLs for exon expression, cis-lncQTLs for lncRNA expression, cis-enQTLs for enhancer expression, and cis-sQTLs for alternative splicing) from 34 tissues in the PigGTEx resource^33^. We performed summary-based heritability enrichment analyses and detected 357 (52.12%) significantly enriched molQTL-trait pairs for 84 out of 147 meta-GWAS summaries (normal test, FDR < 0.05) (Fig. S8f, Table S6). In general, the five types of molQTL were significantly enriched for heritability of all the 15 M_traits (normal test, FDR < 0.05) (Fig. 4c). Furthermore, in muscle (Fig. 4d) and liver (Fig S8g), independent eQTL and sQTL with different ranks explained higher heritability compared to MAF-matched SNPs. These results suggest that variants regulating molecular phenotypes, such as gene expression, play an important role in the genetic mechanism underlying complex traits.

To investigate the relationship between tissue-sharing patterns of eGenes and complex traits, we categorized eGenes into seven tissue-sharing groups^33^. We then performed heritability enrichment analyses for these groups of eGenes using meta-GWAS summary statistics, resulting in 531 significantly enriched gene group-trait pairs for 87 complex traits (normal test, FDR < 0.05) (Table S7). Our enrichment analyses revealed that complex traits were regulated by eGenes with different patterns of tissue-sharing (Fig. 4e). Specifically, we observed a notable enrichment of Backfat thickness (BFT) related traits for eGenes with lower tissue-sharing degree, while ADG-related traits were significantly enriched for eGenes with higher tissue-sharing degree (Fig. 4e). Furthermore, the lead SNP *rs1108824455* for M_BFT acted as an eQTL for *LGALS13* (tissue-specific magnitude = 10) in adipose (*P* = 7.84×10^−23^) (Fig. 4f). *LGALS13* is expressed in lung, duodenum, fetal thymus, jejunum, blood, adipose, ileum, ovary, small intestine, and spleen (Fig. S8h), serving as one of the serum biomarkers in early pregnancy^49,50^. This finding suggests that regulatory variants may affect M_BFT by influencing gene expression in certain tissues during early pregnancy. In addition, the lead SNP *rs324200444* of M_ADG also acted as an eQTL for *ABCC10* in various tissues including the liver (*P* = 2.80×10^−11^), colon (*P* = 7.91×10^−8^), large intestine (*P* = 5.92×10^−7^), and muscle (*P* = 1.34×10^−5^) (Fig.4g). *ABCC10* exhibits widespread expression across different tissues (Fig. S8i) and serves as a genetic marker for pig growth^51^. This finding suggests that regulatory variants affect M_ADG by modulating gene expression in multiple tissues.

### Tissue-specific regulation of GWAS loci

To further investigate the tissue-mediated patterns of genetic regulation of complex traits, we conducted enrichment analyses of GWAS signals using tissue-specific genes across 34 tissues. Gene Ontology (GO) and Kyoto Encyclopedia of Genes and Genomes (KEGG) pathways enrichment analyses of tissue-specific genes confirmed the known biology of respective tissues (Table S8, Fig. S9a). For example, genes highly expressed in muscle were significantly enriched in actin filament binding and muscle contraction (Fig. S9a). Our GWAS signal enrichment analyses demonstrated that significant SNPs of traits were significantly enriched in tissue-specific genes of functionally related tissues (Fig. S9b). For example, the liver was found to be the most enriched tissue for both litter weight (weaning) (M_TLWT_Weaning) (10.14-fold, *P* < 0.001) and body weight (M_BW) (6.84-fold, *P* < 0.001), and the ovary was identified as the most enriched tissue for gestation length (M_GD) (5.62-fold, *P* < 0.001) (Fig. 5a).

**Figure 5.**
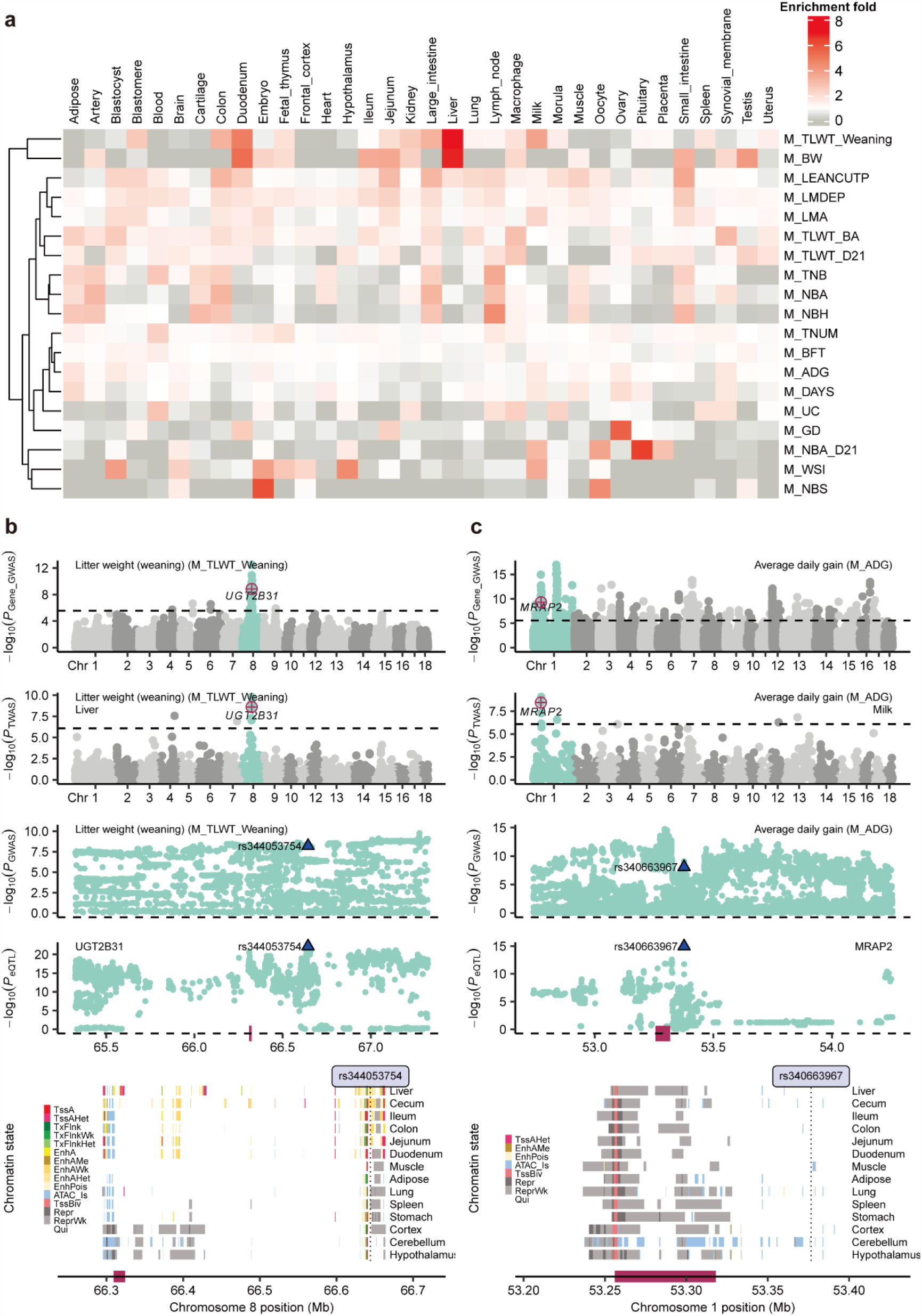
Tissue-specific regulation of GWAS loci. **(a)** Enrichment results for significant associated SNPs of 19 main traits with large sample sizes in tissue-specific functional regions (the top 1,000 tissue-specific highly expressed genes per tissue and their upstream and downstream 100kb regions) in each of 34 tissues. Colors indicate enrichment fold. Rows indicate traits and columns indicate tissues. Enrichment for trait-tissue pairs *E*_T_ = *p*_T_ (proportion of significant SNPs for trait *Tr* located in functional regions of tissue *Ti*)/*q*_T_ (proportion of all SNPs located in functional regions of tissue *Ti*), which calculated by BEDTools v2.25.0^57^. Assoc iated SNPs were resampled 1,000 times with MAF within 0.02 and LD within 0.1 matched to calculate enrichment significance. An *E*_T_ greater than one and *P* less than 0.05 indicates that associated SNPs are significantly enriched in functional regions of tissue *Ti*. (**b**) The association of *UGT2B31* with litter weight (weaning) (M_TLWT_Weaning). The top one Manhattan plot represents the gene-based GWAS results of M_TLWT_Weaning. The top two Manhattan plot represents the single-tissue TWAS results of M_TLWT_Weaning in the liver. Followed by the two following Manhattan plots show the colocalization of M_TLWT_Weaning GWAS (up) and *cis*-eQTL (down) of *UGT2B31* on chromosome 8 in the liver. The blue triangles indicate the colocalized variants of *UGT2B31* in the liver (rs344053754). The bottom panel is for chromatin states around *UGT2B31* on chromosome 8. (**c**) The association of *MRAP2* with average daily gain (M_ADG). The top one Manhattan plot represents the gene-based GWAS results of M_ADG. The top two Manhattan plot represents the single-tissue TWAS results of M_ADG in the milk. Followed by the two following Manhattan plots show the colocalization of M_ADG GWAS (up) and *cis*-eQTL (down) of *MRAP2* on chromosome 1 in the milk. The blue triangles indicate the colocalized variants of *MRAP2* in milk (rs340663967). The bottom panel is for chromatin states around *MRAP2* on chromosome 1.

Furthermore, we integrated multi-omics data from the PigGTEx to explore the detailed regulatory mechanisms of tissue-specific regulation of complex traits. We finally prioritized 16,664 variant-gene-tissue-trait circuits, 19,532 variant-exon-tissue-trait circuits, 3,982 variant-lncRNA-tissue-trait circuits, 3,320 variant-enhancer-tissue-trait circuits, 19,516 variant-splicing-tissue-trait circuits (Table S9). For instance, *UGT2B31*, the most highly expressed gene in the liver compared to other tissues (Fig. S10a), was significantly associated with M_TLWT_Weaning by both gene-based GWAS and TWAS (Fig. 5b). Furthermore, its *cis*-eQTL in the liver colocalized with the GWAS locus (*rs344053754*) of M_TLWT_Weaning (*P* = 5.62×10^−7^), which resides in the active enhancer regions of the liver and intestine but no other tissues (Fig. 5b, Fig. S10b). *UGT2B31* is a metabolic enzyme in the liver of various animals^52–55^. The pattern of *MRAP2* was similar to that of *UGT2B31*, with a higher expression in milk than in most tissues (Fig. S10c). *MRAP2* was also significantly associated with M_ADG in both gene-based GWAS and TWAS. We observed a significant colocalization between the GWAS locus (*rs340663967*) of M_ADG and *cis*-eQTL of *MRAP2* in milk (*P* = 2.86×10^−6^). The colocalized SNP fell into the ATAC region in only muscle and cerebellum (Fig. 5c, Fig. S10d). Interestingly, *MRAP2* was previously identified as a candidate gene for M_ADG in pigs^56^. These results provided important insights that regulatory variants affect gene expression by influencing the activity of regulatory elements in specific tissues, which in turn impact complex traits.

### Gene mapping of complex traits between pigs and humans

To explore the sharing of genetic regulatory mechanisms of complex traits between species, we first conducted heritability enrichment analyses for 169 meta-GWASs in pigs and 136 complex traits in humans based on the orthologues GWAS signals (*P* < 5 × 10^−8^) (Table S10). We obtained 616 significantly enriched pig-human trait pairs (enrichment fold > 1 and *P* < 0.05) (Table S11), including Cholesteryl ester transfer protein activity (S_CEPTA) in pigs *vs*. high cholesterol in humans (enrichment fold = 68.74, *P* = 2.15×10^−2^); Total Cholesterol (S_CHOL_T) in pigs *vs*. Triglycerides in humans (enrichment fold = 18.17, *P* = 1.20×10^−2^), and Feed conversion ratio (30-100kg) (S_FEEDCON_30T100) in pigs *vs*. insulin resistance (HOMA-IR) in humans (enrichment fold = 16.44, *P* = 7.52×10^−3^) (Fig. 6a-b). Permutation analysis demonstrated that the orthologous regions of pig QTLs explained higher heritability of human complex traits compared to randomly selected regions (Fig. S11). We also estimated Pearson’s correlations of pig-human trait pairs based on the absolute *Z*-score of orthologous variants from GWAS summary statistics and found significant correlations of trait pairs with physiological correlations (Fig. 6c, Table S12). For example, the semen sexuality score (S_SESS_DRP) in pigs was significantly correlated with C61 Malignant neoplasm of the prostate in humans (Pearson’s *r* = -0.09, *P* = 2.60×10^−4^) (Fig. 6c). These findings indicate that genetic regulatory mechanisms of certain complex traits were shared between humans and pigs. Furthermore, we discovered that rs322242884 was suggestively associated with L_ADG in Landrace pigs (*Z*-score = -4.53, *P* = 5.80×10^−6^), and its homologous variant rs11877146 was significantly associated with body fat percentage in human (*Z*-score = 6.06, *P* = 1.33×10^−9^) (Fig. 6d). Notably, rs11877146 and rs322242884 were eQTLs for *NPC1* in the muscle of both humans (*P* = 6.70×10^−5^) and pigs (*P* = 7.20×10^−7^), respectively, as well as eQTLs for *TMEM241* in the brain for both species (*P* = 6.70×10^−5^ and *P* = 1.78×10^−4^, respectively), with the similar regulatory effects on gene expression (Fig. 6e-f). Previous studies have linked *NPC1* to body weight and adipocyte processes in a variety of animals^58–61^ and *TMEM241* has been associated with bone degeneration and osteoporosis^62^. These results provided evidence that there might be shared regulatory mechanisms underlying complex traits between humans and pigs.

**Figure 6.**
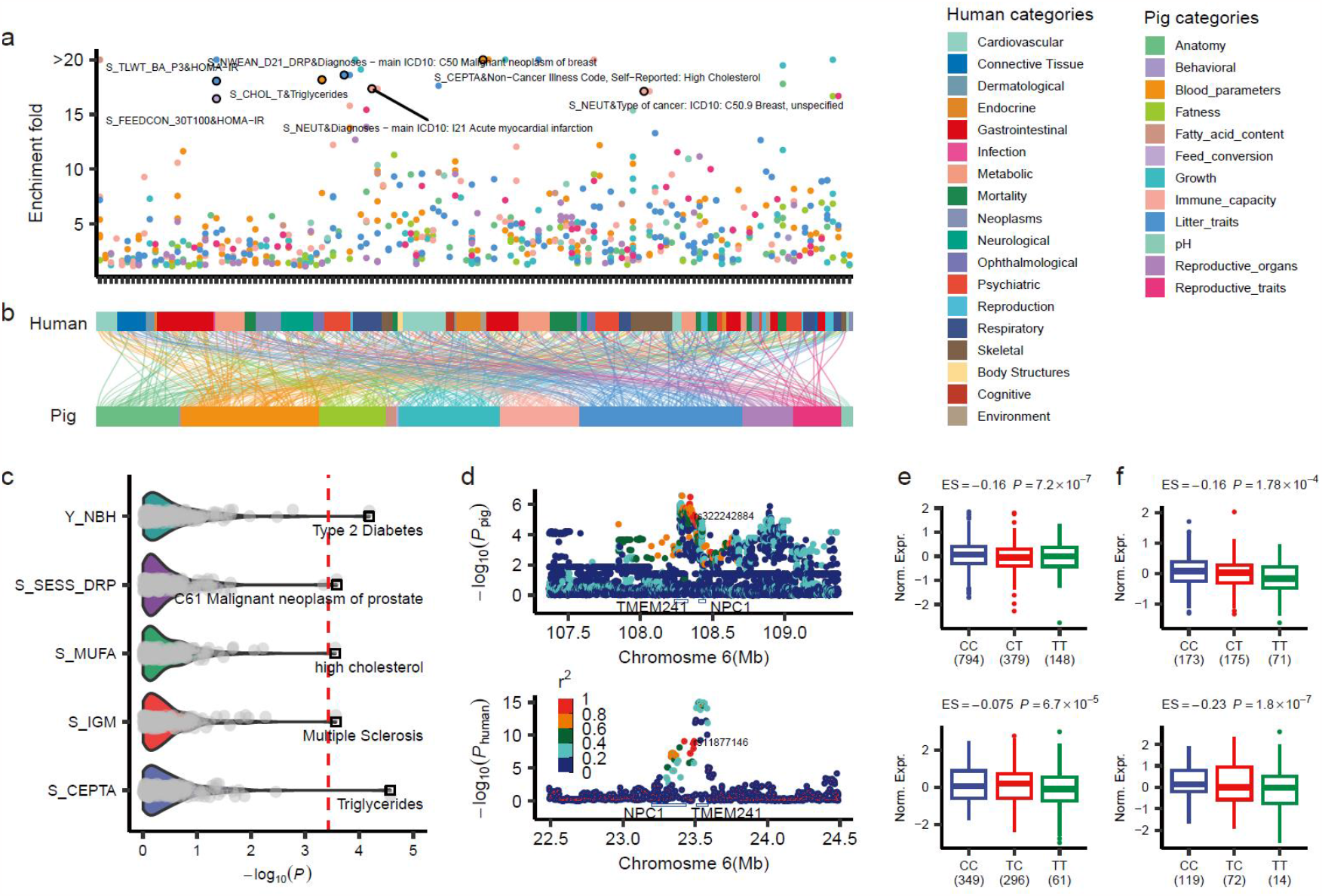
Comparison of complex trait genetics between humans and pigs. (**a)** The heritability enrichment fold between human and pig complex traits is calculated by LDSC. Colors indicate trait categories. (**b**) The alluvium-stratum plot showed the correlation between human and pig complex traits. The alluvium between human-pig trait pairs indicates the heritability enrichment fold > 1 and the *P* < 0.05. Colors indicate trait categories. **(c)** The *P* value was derived from the Pearson’s correlation test of traits between humans and pigs, which was estimated by the absolute Z score of homologous variants from GWAS summary statistics. Each point is a trait pair. The red line is the corrected significant threshold (*P* = 0.05 / 136). Top trait pairs are labeled. **(d-f)** Similar regulatory mechanisms between body fatness rate (BFR) in humans and average daily gain (D_ADG) in pigs. **(d)** The top is a local Manhattan plot of GWAS for D_ADG in pigs. The bottom is a local Manhattan plot of GWAS for BFR in humans. The red triangles represent homozygous variants of humans (rs11877146) and pigs (rs322242884). Colored dots indicate LD with the homozygous variants. **(e)** Top and bottom are the effects of homozygous variants in **(d)** on the expression of homozygous gene *NPC1* in muscle of pigs and humans, respectively. **(f)** Top and bottom are the effects of homologous variants in **(d)** on the expression of the homologous gene *TMEM241* in the brains of pigs and humans, respectively. The significance tests in **(e-f)** were performed by the *wilcox*.*test* function of the *ggsignif* package in R (v4.2.1).

## Discussion

Understanding the genetic foundation of complex traits in pigs have significant implications for improving their genetics, economic contributions, and even medical advancements. While genetic association for major economic traits in commercial pig breeds has been extensively studied, comprehensive GWAS covering a large scale of domesticated pig breeds, as well as a wide range of phenotypes, has not been available. Here, we aimed to establish the largest genetic association atlas of pig complex traits to date by analyzing 70,328 pigs covering 14 pig breeds from various geographic areas.

We identified 6,878 lead variants associated with 139 traits in 169 GWAS meta-analyses (Table S5). The majority of the lead SNPs (99.05%) were located in non-coding regions, including intergenic (39.42%) and intron (43.26%) regions (Fig. S8a). This suggests that these SNPs influence complex traits through playing a crucial role in regulating gene activity^63^. Additionally, the enrichment of lead SNPs in flanking regions of coding sequence further supports the notion that trait-associated SNPs tend to be located in regulatory regions (Fig. 4a and Fig. S8d-e). We also found that lead variants for complex traits usually altered the activity of regulatory elements in specific tissues (Fig. 5b-c). It highlights the importance of tissue-specific gene regulation in determining phenotypic outcomes. The potential epistatic interactions among these genetic variants require further investigation with a larger sample size^64^. The similarity in genetic structure between complex traits in pigs and humans is noteworthy (Fig. 6). This finding suggests that pigs can serve as valuable models for studying human complex traits, offering insights that can contribute to medical advancements in humans.

We make the summary statistics of 268 meta-GWAS available to the research community to facilitate further understanding of the genetic structure of complex traits. Our database provides the comprehensive relationship among genetic variants, genes, tissues and complex traits, which will be useful for dissecting the genetics of complex traits in pigs.

## Acknowledgements

Zhe Zhang acknowledges funding from National Key R&D Program of China (2022YFF1000900), the National Natural Science Foundation of China (32022078), the Local Innovative and Research Teams Project of Guangdong Province (2019BT02N630), and supporting from National Supercomputer Center in Guangzhou, China. Y.C., Zhe Zhang, J.L., X.Liu., S.M., and X.D. acknowledge funding from the China Agriculture Research System (CARS-35). L.F. acknowledges funding from HDR-UK award HDR-9004 and the European Union’s Horizon 2020 research and innovation n program under the Marie Skłodowska-Curie grant agreement No 801215. G.E.L., was supported by USDA NIFA AFRI grant numbers 2019-67015-29321 and 2021-67015-33409 and the appropriated project 8042-31000-112-00-D, “Accelerating Genetic Improvement of Ruminants Through Enhanced Genome Assembly, Annotation, and Selection” of the USDA Agricultural Research Service (ARS). This research used resources provided by the SCINet project of the USDA ARS project number 0500-00093-001-00-D. Mention of trade names or commercial products in this article is solely for the purpose of providing specific information and does not imply recommendation or endorsement by the USDA. The USDA is an equal opportunity provider and employer. L.M. was supported in part by AFRI grant numbers 2020-67015-31398 and 2021-67015-33409 from the United States Department of Agriculture (USDA) National Institute of Food and Agriculture (NIFA). R.X. was supported by Australian Research Council’s Discovery Projects (DP200100499). All the funders had no role in study design, data collection and analysis, decision to publish or preparation of the manuscript.

We thank all the researchers who have contributed to the publicly available data used in this research. For the purpose of open access, the author has applied a Creative Commons Attribution (CC BY) license to any Author Accepted Manuscript version arising from this submission.

## Author contributions

Study design: Zhe Zhang, L.F., F.Z., Y.Z.; Genotype and phenotype data preprocessing: Zhiting Xu, Z.Zhong, H.Zeng, Xiaoqing Wang, L.S., Xue Wang, Y.Wang, Zipeng Zhang, Y.Lin, C.W., J.Z., X.Z., Q.L., J.T., S.D., Yuqiang Liu, X.Pan, X.F., R.L., Z.S., C.C., Q.Zhu.; Genotype imputation: Z.Zhong, Zhiting Xu, H.Zeng, Xiaoqing Wang, H.Y.; Individual GWAS and meta-GWAS analyses: Zhiting Xu, Z.Zhong, B.L., H.Zeng, C.W., Xiaoqing Wang; QTL validation: Q.L., Z.C., Zhiting Xu; GWAS and multi-omics data integration: X.C., Q.L., J.T., J.W., Zhiting Xu; Genetic parameter estimation: Q.L., W.Z., S.S., Y.Wu.; Comparison of pig GWAS with human GWAS: Q.L., Y.Wu., Z.B.; Critical interpretation of analytical results: L.F., Zhe Zhang, B.L., J.T., Y.G., G.E.L., P.K.M., M.F., S.L., F.Z, Y.Z., Q.Zhang, G.E.L., X.S., R.X., L.M., M.S.L., G.Sahana, G.Su, Yang Liu, P.L.; Contribution of data and computational resources: Zhe Zhang, L.F., F.Z., Y.Z., Q.Zhang, Y.C., J.L., X.D., X.Li, M.L., G.T., M.F., P.K.M., A.C., M.A., D.C.P., M.B., R.H., P.L., X.Y., H.Zhang, X.Liu, D.M., Y.P., Q.W., K.L., L.B., Z.Zhou, Zhong Xu, X.Peng, S.M.; Drafting the manuscript: Zhe Zhang, L.F., Zhiting Xu, Q.L.. All authors read, edited, and approved the final manuscript.

## Methods

### Ethics

This is not applicable because no biological samples were collected, and no animal handling was performed for this study.

### GWAS dataset

In total, we collected 70,328 pigs with genotype and phenotype data from 59 study populations (14 public populations) covering 14 pig breeds (Table S1). We conducted comprehensive data preparation and standardization for the study data regarding phenotype and genotype according to the previously published protocol^65^.

### Genotype data

We genotyped these pigs from these 59 populations using low-coverage sequence (N = 2,873) or genotyping arrays, including the Illumina Porcine SNP60K Bead Chip (N = 10,870), the GeneSeek Genomic Profiler (GGP) Porcine SNP80 BeadChip (N = 4,724), the GGP Procine SNP50 BeadChip (N = 29,789), the KPS Porcine Breeding Chip (N = 21,618), the GenoBaits Porcine SNP50K BeadChip (N = 454). We constructed a standard pipeline to uniformly process individual-level genotype data for all 59 populations. Briefly, we first converted the coordinate of the genomic version of genotype data to the Sscrofa11.1 (v100) and only kept the autosomal biallelic SNPs. To remove the outliers within each population, we performed principal component analysis (PCA) for each of the 59 populations using PLINK (v1.9)^34^ based on LD-independent SNPs with parameter: “*--mind 0*.*1 --geno 0*.*9 --maf 0*.*01 --indep-pairwise 50 5 0*.*5 -- pca 10*”. We visualized the principal components (PCs) of each population in R (v3.4.3) and then excluded a total of 1,086 individuals who were outliers using PLINK (v1.9). Finally, we retained 69,242 individuals for downstream analyses, including 20,706 Duroc pigs, 34,540 Yorkshire pigs, 9,159 Landrace pigs and 4,837 other pigs.

### Genotype imputation

To obtain genotype data at whole-genome sequence (WGS) level, we performed genotype imputation for each population based on multi-breed Pig Genomics Reference Panel (PGRP v1) from PigGTEx^33^, which consists of 42,523,218 autosomal biallelic SNPs from 1,602 WGS samples covering over 100 pig breeds. We firstly removed duplicate alleles from array data using PLINK (v1.9)^34^ with parameter: “*--list-duplicate-vars ids-only suppress-first, --exclude plink*.*dupvar --recode vcf bgz*” and kept biallelic SNPs using BCFtools (v1.9)^57^. We then employed conform-gt program (http://faculty.washington.edu/browning/conform-gt.html) to revise strand inconsistencies of SNPs based on pre-phasing genotype data^66^. We imputed the genotype data of target populations to sequence level using Beagle (v5.1)^67^ and filtered out variants with dosage R-squared (DR2) < 0.8 and MAF < 0.01 within each population. Finally, we retained a total of 28,297,603 SNPs across all 59 populations for downstream analysis (Table S1).

To evaluate the accuracy of genotype imputation, we employed two strategies (Fig. S1a). (1) We conducted 20 rounds of five-fold cross-validation using genotype data or 60,720 samples from 53 GWAS populations that had individual-level genotype data. Specifically, in each round of cross-validation, we randomly selected 20% of SNPs in chromosome 6 of the target panel as a validation set and imputed them using PGRP as a reference panel via Beagle (v5.1). We measured the imputation accuracy by calculating the concordance rate and Pearson’s correlation between the imputed and true genotypes in the validation set. (2) We obtained 65 WGS samples from NCBI that were independent of PGRP, comprising of 35 Duroc pigs (PRJNA712489) and 30 Suhuai pigs (PRJNA791712) (Table S2). We employed Trimmomatic (v0.39)^68^ to filter out the adaptors and low-quality reads, mapped clean reads to Sus scrofa11.1 (v100) using BWA-MEM (v0.7.5a-r405) with default parameters^69^, and marked duplicated reads using Picard (v2.21.2) (http://broadinstitute.github.io/picard/). We called SNPs for these samples using Genome Analysis Toolkit (GATK) (v4.1.4.1)^70^ with parameter: “*QD> 2, MQ < 40, FS > 60, SOR > 3, MQRankSum < -12*.*5* and *ReadPosRankSum < -8*”, resulting in 17,182,138 and 15,696,890 biallelic autosomal SNPs for Duroc and Suhuai, respectively. For the purpose of evaluating the accuracy of genotype imputation, we masked SNPs that were not overlapped with these SNPs obtained from SNP array and then imputed them to WGS level using PGRP as reference panel via Beagle (v5.1). Finally, we calculated the concordance rate and Pearson’s correlation between imputed genotypes with DR2 > 0.8 and MAF > 0.01 and those called directly from the WGS.

### Phenotype data

A total of 286 complex traits (15 binary traits and 271 continuous traits) were available for the 59 populations (Table S3), which belonged to five main trait-categories (i.e., Reproduction, Meat and Carcass, Health, Production, and Exterior) and 17 sub trait-categories (i.e., Litter, Reproductive, Growth, Reproductive organs, Blood parameters, Immune capacity, Anatomy, Fatness, Fatty acid content, Feed conversion, Conformation, Meat color, Chemistry, Feed intake, pH, Texture, and Behavioral).

In particular, 49 out of 286 traits have phenotypic records in multiple time points for the same individual (e.g., sperm traits and litter sizes, detailed in Table S3) and were referred to as “multiple time points trait” (i.e., MT-trait). For these 49 MT_traits, we calculated the deregressed proofs (DRP) as their phenotype measures using DMU (v6-R5-2-EM64T)^71^. We first estimated breeding values (EBV) in each population based on pedigree information using a single-trait repeatability model implemented in the DMUAI module of DMU. The statistical model is:

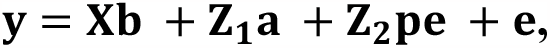

where **y** is the vector of phenotypic values for all individuals; **b** is the vector of the effects of covariates (e.g., year-season of ejaculation, age of pigs at months or collection interval (days)); 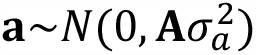 is the vector of additive genetic effects, with **A** and 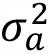 denoting the pedigree-based additive genetic relationship matrix and additive genetic variance; 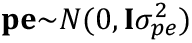 is the vector of permanent environmental effects with 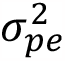 denoting the identity matrix and the permanent environmental variance; **X, Z**_**1**_, and **Z**_**2**_ are the incidence matrices assigning observations to covariates effects, additive genetic effects, and permanent environmental effects, respectively; 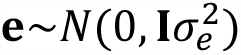 is the vector of random residual effects, with **I** and 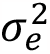 denoting the identity matrix and the residual variance. To eliminate the bias from relatives, we calculated the DRP and weights for each pig using the methods described by Garrick et al.^72^ with the following model:

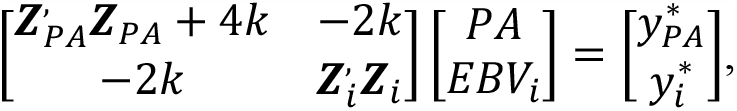

where 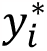 is information equivalent to a right-hand-side element pertaining to the individual, *PA* is the parental average EBV; EBV_i_ is the EBV for animal *i*; 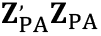 and 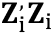 reflect the unknown information content of the parental average and individual (plus information from any of its offspring and/or subsequent generations). Their formulas were as follows:

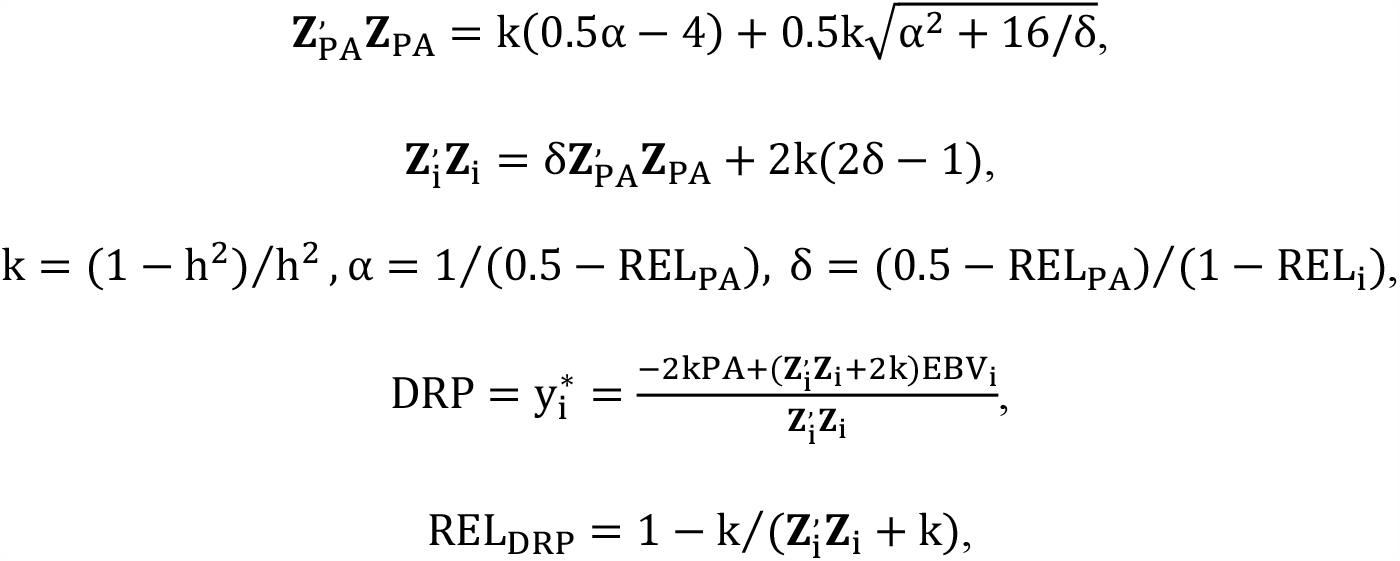

where *h*^2^ is the estimated heritability; *REL*_*PA*_ is the reliability of the parental average EBV; *REL*_*i*_ is the reliability of the EBV for animal *i*; REL_DRP_ is the reliability of the DRP for animal *i*. The weights can be derived from 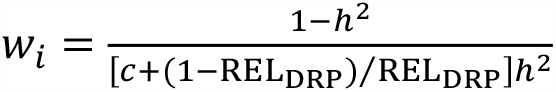, where c= 0.2 is assumed to represent the proportion of genetic variation for which genotypes cannot account is 0.2. Finally, we used the DRP and weights for each pig above for the downstream association analysis.

### Individual GWAS

We conducted individual GWAS for each trait in each population as described below and referred to this as “individual GWAS” throughout the manuscript (Table S3).

For binary traits, we performed association analysis with a logistic mixed model using fastGWA-GLMM implemented in GCTA (v1.94.0)^73^. The statistical model is:

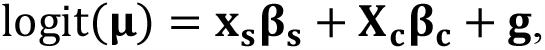

where **y** is a vector of phenotypic values, **μ** is a vector of μ_i_= P(y_i_= 1|x_si_, X_ci_, g_i_) with *μ*_*i*_ being the probability of subject *i* being a case given the subject’s genotype *x*_*si*_, covariates *X*_*ci*_ and random genetic effect *g*_*i*_; **x**_**s**_ is a vector of genotype variables of a variant of interest with its effect β_s_ ; X_c_ is the incidence matrix of fixed-effect covariates (farms, sex, year, season and the first five genotype PCs) with their corresponding coefficients β_c_; *g* is a vector of effects that capture genetic and common environmental effects shared among related individuals, 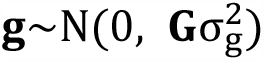 with **G** being the sparse GRM with all the small off-diagonal elements (for example, those <0.05) set to zero and 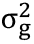 being the corresponding variance component.

For quantitative traits, we performed association analysis with a mixed linear model using fastGWA implemented in GCTA (v1.94.0)^74^. The statistical model is:

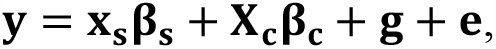

where **y, x**_**s**_, **β**_**s**_, **X**_**c**_, **β**_**c**_ and **g** are the same as those in the above logistic mixed model; *e* is the vector of residuals with 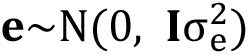.

Specifically, for 49 MT-traits, we employed MMAP (v2021_08_19_22_30.intel) (https://mmap.github.io/) to perform association analysis based on their DRP and weights for each pig. We conducted the individual GWAS based on a mixed linear model:

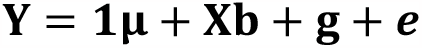

where **y** is the vector of DRP for the given trait, **μ** is the global mean, and **1** is a vector of ones; **X** is the genotype of a candidate variant (coded as 0, 1, or 2 copies of the minor allele) for the animals with observations in **y**, and **b** is a vector of marker effects; **g** is a vector of polygenic effects accounting for population structure with 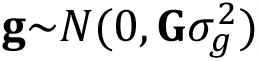, where the genomic relationship matrix (**G**) was built using the imputed SNPs and 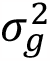 is the genetic variance, and **e** is a vector of random residual errors with 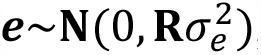, where 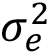 is residual error variance and **R** is a diagonal matrix that adjusts 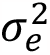 to account for the heterogeneous variance of DRP for each pig, the weights included in **R**.

### Meta-analysis of GWAS

To enable individual GWASs from different populations to be comparable in the meta-analysis, we checked all summary statistics based on EasyQC^65^.

First, to detect issues related to trait transformations, we first examined the relationship between the inverse of the median standard error of all SNPs beta estimates and the square root of the sample size (SE-N plot) across multiple study files for each trait. For outliers, we examined the raw phenotype data and reran association analysis. The calibration factor c of the SE-N plot was approximated from the autosomal SNPs of the PGRP reference panel as 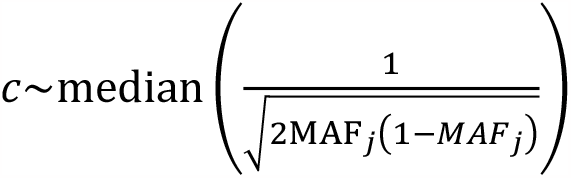. Second, we examined the analytical issues for each study by comparing the reported *P* values of each SNP with the *P* values computed from the *Z*-statistics (*Z*-statistics = *β*_*j*_/*SE*(*β*)_*j*_) based on reported beta estimate and standard error (P-Z plots). Third, we plotted the effect allele frequency (EAF) from study-specific against EAF from PGRP to identify strand issues or allele miscoding that could severely reduce statistical power. Fourth, we grasped the potential problems with population stratification by the genomic control (GC) inflation factor (λ_GC_, from 0.86 to 2.39 with an average of 1.11). After we reconstructed the association analyses by using the first five principal components as additional covariates, the λ_GC_ decreased (from 0.56 to 1.58 with an average of 1.04). Fifth, we excluded SNPs with missing or nonsensical information (e.g., *P* values < 0 or >1, or non-numeric values such as “NA”) from summary statistics.

We performed meta-analyses on the cleaned GWAS results for each trait using METAL (v2011-03-25)^75^, based on an inverse variance-weighted fixed effects model that weights effect size estimates according to estimated standard errors and allows for different population frequencies of genotypes and alleles. Genomic control correction was applied for all input files in the analysis. SNPs included in the meta-analysis were present in at least one individual GWAS, and the total number of SNPs for each trait is shown in Table S4.

### Definition of QTL

For both individual GWASs and meta-GWAS, we used *P* < 5.0 × 10^−8^ as the genome-wide significance threshold and defined lead SNPs and QTLs on the basis of genomic position. For each GWAS summary, we defined the significant SNP with the smallest *P*-value in each chromosome as the first lead SNP and the significant SNP with the smallest *P-*value outside the 1Mb-region upstream and downstream of the first lead SNP as the second lead SNP. This process was iterated until no significant SNPs were left in that chromosome. Different traits can share lead SNPs. We defined the two most distant significant SNPs within 0.5Mb on each side of the lead SNPs as the boundaries of the QTLs. In addition, we performed a stepwise conditional analysis to extend the candidate regions and define broad QTLs, in which adjacent significant SNPs within the broad QTL region are within 1Mb apart.

### QTL Validation

To validate the QTL regions we identified, we used the following three strategies.

First, we compared the QTL regions with those for the same traits reported in the Pig Quantitative Trait Locus (QTL) database (Pig QTLdb version 46)^23^ (https://www.animalgenome.org/cgi-bin/QTLdb/SS/index). We performed a filtering process on the downloaded QTL regions, excluding those with missing start/end position information and those smaller than 1bp or larger than 1Mp. This resulted in a final retention of 302,784 autosomal QTL regions. Among these, we successfully matched 151 traits with traits in our study. Regions, where there was at least 1bp of overlap with the same trait, were considered successfully validated (defined as ‘TRUE’), while those without overlap were classified as ‘FALSE’.

Second, we validated the QTL regions in independent populations. For this, we performed nine individual GWASs for average daily gain (ADG) on a total of 42,790 pigs, 88,984 pigs, and 69,606 pigs from three populations of Duroc, Landrace, and Yorkshire, respectively, using the MLMA model of GCTA (v1.94.0)^76^. This statistical model was consistent with one of the fastGWA models used in our study. Subsequently, we conducted within-breed meta-GWAS analyses and all-breed meta-analysis using the same method as in this study. We identified the QTL regions in these meta-analyses using the same method as in our study and calculated the enrichment fold of these regions in the QTL regions of ADG detected in our study.

Third, we used information on suggestively significant lead SNPs (*P* < 1.0 × 10^−5^) for breed-level genomic prediction to validate the functional reliability of the QTLs. For this, we performed genomic predictions in seven pig breeds from PGRP, including 54 Meishan, 24 Erhualian, 41 Jiaxinghei, 226 Yorkshire, 51 Landrace, 138 Duroc, and 43 Pietrain pigs. We extracted the genotypes of the lead SNPs from PGRP using Bcftools (v1.9)^57^ and their effect sizes from GWAS summary statistics. Whereafter, we used a linear mixed model to fit the genotype and effect size for genomic prediction in each breed. The model formula we used for each breed was:

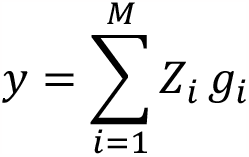

where *y* is a vector of predicted phenotypes, *g*_*i*_ is the effect size of lead SNP *i* in GWAS summary statistics, *Z*_*i*_ is the vector of the genotype of lead SNP *i* containing 0, 1 and 2. We fitted the model using R v 4.2.1.

### Pleiotropic variants across breeds and traits

To identify variants with effects on traits shared among breeds, we extracted the effect sizes and standard errors of lead SNPs from a total of 36 meta-analyses for 12 traits in Duroc, Landrace and Yorkshire pigs. We then used METASOFT (v2.0.1)^44^, a procedure that corrects for the effect of sample size on association analysis, to calculate the posterior probability of the lead SNP effect for each trait in each breed. We employed the same method to identify GWAS variants with pleiotropic effects on multiple traits. We considered an M-value greater than 0.9 as evidence of an effect.

### Annotation and enrichment of significant/lead variants in functional categories

To investigate the molecular mechanisms of significant/lead SNPs, we examined multiple layers of biological data.

First, we annotated significant/lead SNPs in several genomic categories: (i) 20 genomic variants, including intron variants and intergenic region variants, using SnpEff (v.4.3)^77^. (ii) the seven groups categorized by genomic locations with respect to protein-coding genes, i.e., CDS, promoter (100kb upstream and downstream of the protein-coding gene TSS), 5’UTR + 2kb upstream, 3’UTR + 2kb downstream, protein-coding genes, non-protein-coding genes, and intron regions. (iii) the downloaded mammalian conserved elements identified from Multiple Sequence Alignments (MSA) using the Genomic Evolutionary Rate Profiling (GERP) software based on 103 mammals (https://ftp.ensembl.org/pub/release-100/bed/ensembl-compara/103_mammals.gerp_constrained_element/). (iv) the 14 chromatin states detected from 14 major pig tissues^78^ to investigate the regulatory function. (v) the tissue-specific functional regions of 34 tissues in FarmGTEx^33^. Here, we borrowed the top 1,000 tissue-specific highly expressed genes, along with their upstream and downstream 100kb regions in each tissue, to represent the tissue-specific functional regions.

Second, we estimated the enrichment and significant *P*-value of significant/lead SNPs across the various genomic categories. For genomic variants, we used the *oddsratio* function of fmsb (v0.7.5) package^79^ in R (v4.1.2) to perform enrichment and estimate significance. The enrichment for category *C* (*E*_C_) = *p*_C_ (proportion of significant/lead SNPs located in category *C*) / *q*_C_ (proportion of all SNPs located in category *C*). For the genomic regions of protein-coding genes, conserved elements, chromatin states and tissue-specific functional regions, we used two methods to estimate the enrichment: (i) we employed the R/Bioconductor package locus overlap analysis (LOLA v1.22.0)^80^ to estimate the enrichment and *P*-values, and (ii) we used BEDTools (v2.25.0)^57^ to estimate the enrichment. The enrichment for category *C* (*E*_C_) = *p*_C_ (proportion of category *C* in all significantly enriched trait-category pairs *E*_T_) / *q*_C_ (proportion of category *C* in the genome). Here, the enrichment for trait-category pairs *E*_T_ = *p*_T_ (proportion of significant/lead SNPs for trait *T* located in category *C*)/*q*_T_ (proportion of all SNPs located in category *C*). We performed a permutation test by resampling the association signals 10,000 times to determine if the observed SNPs located in the annotation category were greater than expected by chance, using the R package regioneR (v1.24.0)^81^. Additionally, we resampled the SNPs matching the MAF (within 0.02) and LD (within 0.1) of the association signals 1,000 times for the permutation test. An *E*_C_ greater than one and a *P*-value less than 0.05 indicated that significant/lead SNPs were significantly enriched in category *C*.

In addition, to understand the evolutionary sequence conservation of association variants, we downloaded PhastCons scores for 100 vertebrate species from UCSC (http://hgdownload.cse.ucsc.edu/goldenpath/hg38/phastCons100way/hg38.100way.phastCons/). We converted the Wiggle files of PhastCons scores to BED files using the BEDOPS tool (v2.4.40)^82^, and then we lifted them over from the human genome 38 (h38) to Sscrofa11.1 using UCSC’s LiftOver tool^83^.

### Summary-based genetic parameter estimation

To estimate the genetic parameters for all pig complex traits, we first harmonized all 268 GWAS summary data using the *munge_sumstats*.*py* function of the linkage disequilibrium score regression (LDSC v1.0.1)^43^ with parameters: “--sumstats --N –out”, and estimated linkage disequilibrium (LD) score from PGRP using PLINK (v1.90)^34^ with parameter: “--ld-wind-kb 1000”. Second, we estimated the narrow-sense heritability for complex traits based on summary using LDSC (v1.0.1) with parameters: “--h2, --ref-ld-chr, --w-ld-chr and --out”.

### Heritability enrichment of regulatory variants

To investigate the impact of regulatory variants on complex traits, we extracted significant *cis*-molQTLs from five molecular phenotypes in 34 tissues, including 2,930,627 *cis*-eQTLs for protein-coding gene expression, 2,842,703 *cis*-eeQTLs for exon expression, 2,628,257 *cis*-sQTLs for alternative splicing, 2,703,774 *cis*-enQTLs for enhancer and 2,056,718 *cis*-lncQTLs for lncRNA^33^. We used the BLD-Thin model of LDAK (v5.0)^84^ to estimate the heritability and performed heritability enrichment analysis for these molQTLs in 169 meta-GWAS summaries that detected lead SNPs using an optional parameter: “-*-check-sums NO*”.

To further explore the effects of independent regulatory variants on complex traits, we divided independent eQTLs into three groups based on their rank for each eGene from muscle and liver tissues, including primary-, secondary- and third-rank independent eQTLs. We performed the same heritability enrichment analysis for these independent eQTLs in the 169 meta-GWAS summaries. We did not consider the results for enrichment folds less than 0. Additionally, we obtained *P*-values based on the enrichment fold and their standard errors using a one-sided normality test. We adjusted the *P*-value using the *p*.*adjust* function with the FDR method in R v4.2.1. The heritability enrichment with FDR < 0.05 was considered a significant pair. The formula for estimating the *P*-value was:

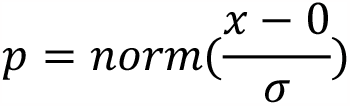

Where *x* represented the heritability enrichment fold, and *σ* was the standard error of the heritability enrichment fold.

To evaluate the performance of heritability enrichment for independent molQTLs, we randomly selected the same number of MAF-matched SNPs for each rank of independent molQTLs for muscle and liver tissue and performed heritability enrichment. We extracted the total SNPs heritability contributed by each category to compare the performance of heritability enrichment.

### Heritability enrichment of tissue-sharing/specific genes on complex traits

To investigate the regulatory patterns of tissue-sharing/specific genes for complex traits, we conducted heritability enrichment analysis of these genes in GWAS summaries using LDAK (the BLD-Thin model)^84^. Initially, we categorized protein-coding genes into seven groups (1-5, 6-10, 11-15, 16-20, 21-25, 26-30, 31-34 tissues) based on the magnitude of tissue-sharing/specificity derived from the *cis*-eQTL meta-analysis results across all 34 tissues^33^. Subsequently, we extracted significant *cis*-eQTLs for each gene and organized them according to their respective tissue-sharing/specific gene groups. Next, we randomly selected 500,000 variants for each gene group to generate an annotation file. Finally, we used the annotation file to calculate the tagging file and conducted the heritability enrichment analysis.

### Colocalization of GWAS summary with cis-molQTL

To investigate the contribution of molecular phenotypes to the genetic regulation of complex traits, we performed a colocalization analysis of molQTL and GWAS signals using fastENLOC (v1.0)^85^. The details of our colocalization methods have been described in our previous work^33^.

### Summary-based transcriptome-wide association study (TWAS)

To explore whether the overall *cis*-genetic component of a molecular phenotype is associated with complex traits, we conducted both single- and multi-tissue TWAS using S-PrediXcan^86^ and S-MultiXcan in MetaXcan (v0.6.11)^87^, based on summary statistics from meta-GWASs. Our TWAS methods have been previously described in detail^33^. We applied the Bonferroni correction for multiple testing and considered a corrected *P*-value of less than 0.05 to be significant.

### Mendelian randomization (MR) analysis between molQTL and GWAS loci

To infer the causality between molecular phenotypes and complex traits, we conducted an integrative MR analysis using the SMR tool (v1.03) with genetic variants as instrumental variables^88^. The method used has been previously described in detail^33^. To account for multiple testing, we applied the Bonferroni correction and defined a corrected *P*-value of less than 0.05 as significant.

Finally, we prioritized variant-gene/exon/lncRNA/enhancer/splicing-tissue-trait circuits that were validated by at least one method, including TWAS, colocalization, and MR. These circuits exhibited significant tissue-trait associations in enrichment analyses of GWAS significant signals and tissue-specific functional regions.

### Heritability enrichment of human complex traits

To investigate whether the regulatory mechanisms of complex traits were conserved between humans and pigs, we used lead variants with extended windows in 169 pig complex traits to determine the heritability enrichment in human complex traits. Initially, we obtained public GWAS summary statistics for 136 human complex traits, representing 18 trait domains (Table S10). We then mapped the genomic regions located 1 Mb upstream and downstream of the lead variants for each pig complex trait to the human genome (GRCh38/hg38) using UCSC’s LiftOver tool^83^. Subsequently, we implemented heritability enrichment analysis using these genomic regions for the 136 human complex traits by LDSC (v1.0.1)^43^. Finally, we selected human-pig trait pairs with a heritability enrichment fold greater than 1 and a *P*-value less than 0.05 for further downstream analysis.

We also conducted a validation study to evaluate the performance of heritability enrichment of pig QTL regions in humans. For this, we randomly selected QTL regions and performed heritability enrichment analysis. Initially, we removed the regions already mapped with pig QTL regions based on human genome information. Next, we randomly selected an equal number of regions with matching widths from the remaining human genome for each pig complex trait. Finally, we used these selected regions for heritability enrichment analysis on the summary statistics of 136 human complex traits.

### The correlation between humans and pigs in GWAS summary statistics

To explore the correlation between pigs and humans in GWAS summary statistics, we first obtained homozygous variants shared between pigs (version: Sus scrofa11.1) and human (GRCh38/hg38) using LiftOver^83^. Second, we matched the homozygous variants for 268 pig GWAS summary statistics and 136 human GWAS summary statistics. Third, we performed a Pearson correlation test between the absolute value of the Z-score of homozygous variants from humans and pigs in R v4.2.1. We considered a threshold of 0.05/136=3.68×10^−4^ as significant for the trait pairs.

### Reporting summary

Further information on research design is available in the Nature Research Reporting Summary linked to this article.

